# Gelatinase B/matrix metalloproteinase-9 is a phase-specific effector molecule, independent from Fas, in experimental autoimmune encephalomyelitis

**DOI:** 10.1101/321661

**Authors:** Estefania Ugarte-Berzal, Nele Berghmans, Lise Boon, Erik Martens, Jennifer Vandooren, Bénédicte Cauwe, Greet Thijs, Paul Proost, Jo Van Damme, Ghislain Opdenakker

## Abstract

Gelatinase B/matrix metalloproteinase-9 (MMP-9) triggers multiple sclerosis (MS) and the animal model of experimental autoimmune encephalomyelitis (EAE) by the breakdown of the blood-brain barrier. Interestingly, MMP-9 is beneficial in systemic autoimmunity caused by Fas-deficiency. Fas-deficient (*fas^lpr^)* and Fas-ligand-deficient mice are protected against EAE. We here investigated the interaction between Fas and MMP-9 in the setting of induction of EAE and compared short- and long-term effects. We provoked EAE with myelin oligodendrocyte glycoprotein (MOG) peptide and compared EAE development in four genotypes (wild-type (WT), single knockout *mmp*-*9*^−/−^, *fas^lpr^*, and *mmp-9*^−/−^*/fas^lpr^*) and monitored leukocytes, cytokines and chemokines as immunological parameters. As expected, *fas^lpr^* mice were resistant against EAE induction, whereas MMP-9 single knockout mice were not. In the double *mmp-9 ^−/−^/fas^lpr^* mice the effects on disease scores pointed to independent rather than interrelated disease mechanisms. On a short term, leukocytes infiltrated into the brain and cytokines and chemokines after EAE induction were significantly higher in all the four genotypes studied, even in the *fas^lpr^* and *mmp9-/-/f* as ^*lpr*^, which did not develop clinical disease. The levels of MMP-9 but not of MMP-2 were increased in the brain and in the peripheral organs after EAE induction. After 40 days all the animals recovered and did not show signs of EAE. However, the absence of MMP-9 in the remission phase suggested a protective role of MMP-9 in the late phase of the disease, thus single *mmp-9* ^−/−^ mice presented a delayed onset and remission in comparison with WT animals suggesting a phase-dependent role of MMP-9 in the disease. Nevertheless, the levels of some cytokines and chemokines were remained higher than in control animals even 100 days after EAE induction, attesting to a prolonged state of immune activation. We thus yielded new insights and useful markers to monitor this activated immune status. Furthermore, MMP-9 but not MMP-2 levels remained increased in the brains and, to a higher extend, in the spleens of the WT mice even during the remission phase, which is in line with the role of MMP-9 as a useful marker and a protective factor for EAE in the remission phase.

## 1. Introduction

Multiple sclerosis (MS) is a devastating autoimmune disease in need of alternative pathophysiological insights and new pharmacological targets [1,2]. Despite major efforts, direct targeting of the adaptive arm of immunity by eliminating T or B cell clones has failed so far for MS treatment, whereas innate immune molecules, including interferons for substitution therapy, minocycline as protease inhibitor [3, 4, 5] and cell adhesion molecules as disease targets, are yielding gradually improved therapeutic schemes. The replacement of injectable drugs by oral ones is a major step forward [5,6]. Specifically, oral and inexpensive minocycline can be used when other drugs are excluded based on prescription and insurance requirements [5]. Targeting adaptive immune mechanisms indirectly with oral fingolimod, which blocks T and B cell egress, shows a better outcome for patients than parenteral interferon [6]. These examples illustrate that gradual improvements towards compliant therapies are possible and that the study of disease mechanisms remains essential [7].

The definition of genetic factors and their interactions in MS and animal models of experimental autoimmune encephalomyelitis (EAE) may give clues for the development of better diagnostics and therapeutics. One gene that has been relatively neglected so far as a disease marker or therapeutic target encodes the apoptosis-inducing factors of the Fas system. Nevertheless, solid evidence exists that both Fas ligand (FasL/CD95L) as well as Fas receptor (Fas/CD95) are critical factors in determining the outcome of antigen-specific EAE [8,9]. Indeed, fas- and fasL-deficient mice are resistant to develop EAE. In addition, inflammatory lesions in *fas*-deficient *lpr* mice (*fas^lpr^*) contain less apoptotic cells than those observed in wild-type lesions. Furthermore, in cell transfer experiments, Fas expression in recipient animals is important for EAE progression because *fas^lpr^* recipients still develop lower disease scores despite transfer of *fas^lpr^* or wild-type T lymphocytes. However, other factors, such as (auto)antigen generation and presentation, are necessary for EAE disease to occur and develop, because T cell receptor (TCR) transgenic mice for myelin antigen on an *lpr* background still produce EAE [10].

A natural way to generate autoantigens for MS and EAE is by extracellular proteolysis or other posttranslational protein modifications in an inflammatory context. Such extracellular proteolysis may be provided by host proteases such as MMP-9, abundantly induced by cytokines and chemokines in inflammation [11]. The focus on proteolysis in MS has been primarily on studies of the inducible matrix metalloproteinase-9 (MMP-9) [12]. This enzyme was first associated with MS [13]. Young mmp-9 knockout mice show resistance against EAE [14] and MMP inhibitors and double mmp-2/mmp-9 deficiency were found to be protective, also in adult mice [15,16]. Recently, targeting MMPs with minocycline as the most potent tetracycline inhibitor of MMP-9 [4], has yielded promising results in MS patients under monotherapy [5]. However, although MMP-9 activity at the blood-brain barrier has recently been shown to constitute a critical event in early MS lesion development [17], the MMP-9 gene has not been found as a disease susceptibility factor in genetic screenings for MS [18]. As a contrast, instead of a disease-causing function in organ-specific autoimmunity [19], a protective role has been described for MMP-9 in systemic autoimmune disease and also this role seems to depend on proteolytic antigen processing [20]. Indeed, when the MMP-9 gene was knocked out in fas-deficient lpr mice (*fas* ^lpr^), that are prone to develop systemic autoimmune disease, the animals were not resistant, but became instead more susceptible to develop lupus-like disease. In this phenotype, autoantibodies are generated against ubiquitous intracellular proteins, many of which are substrates of MMP-9 [20, 21].

Another often underestimated aspect when using EAE models is the duration of the experiments. Most often, EAE experiments are done over a time interval of only a few weeks. This interval is sufficient and usually employed to demonstrate beneficial aspects of gene knockouts or novel treatment schemes. However, and as we experienced in preliminary experiments studying therapeutic effects, gene knockout exploration and analysis of known drugs may fade out on longer duration.

We here addressed the questions (i) whether the “spontaneous” clinical effects of deletion of Fas (fas^lpr^) and MMP-9 (mmp-9^−/−^), as previously observed [20], are similar or different after induction of autoimmunity with a known autoantigen and whether these are additive, synergistic or antagonistic (ii) which of both molecules, Fas and MMP-9, drives clinical outcome in the setting of an organ-specific autoimmune reaction (iii) which cellular and molecular mechanisms are behind the clinical outcomes and how can these be used in monitoring disease states and (iv) what is the long-term progression of EAE and the evolution of immune cells and molecules.

## 2. Materials and methods

### 2.1 Generation and maintenance of mice

To exclude confounding genetic influences of the genetic background we extensively backcrossed our original mouse line [14]. In the 10^th^ backcross generation of MMP-9 knockout mice into C57/Bl6, the mice retained brown fur [20], suggesting that a dominant hair color determinant is located near the mouse MMP-9 gene. In addition, our knockout line has a severe subfertility phenotype [22]. For these reasons and in view of the increasing awareness about possible confounding influences imposed by differences in genetic backgrounds between wild-type and knockout mice [23,24] and by possible interference by differences in environmental conditions [25] and to avoid clinical bias on the basis of fur color, we further backcrossed our *mmp-9* knockout mice till these became black [26]. We used in all forthcoming experiments black animals from the 13^th^ generation backcross into C57Bl/6. All mice were bred in specific pathogen-free (spf) insulators at the Rega Institute for Medical Research and under exactly the same environmental conditions (e.g. food, day night cycle, bedding) for all genotypes. At regular time intervals, we screened for the presence of a panel of common mouse microbes as a control for absence of infections. Induction and follow-up of EAE evolution were carried out with adult (8–10 week old) male and female mice under SPF housing conditions. During this period, the mice received appropriate nutrition and acidified drinking water without antibiotics. All procedures were conducted in accordance with protocols approved by the local Ethics Committee (Licence number LA1210243, Belgium).

### 2.2. Reagents

Mycobacterium tuberculosis strain H37Ra, Incomplete Freund’s Adjuvant (IFA) and Complete Freund’s Adjuvant (CFA) were purchased from Difco Laboratories (Detroit, MI, USA). Pertussis toxin was purchased from List Biological Laboratories (Campbell, CA, USA). Myelin oligodendrocyte glycoprotein peptide (MOG35–55) was produced by Fmoc (fluorenylmethoxycarbonyl) solid phase peptide synthesis, purified by reversed phase chromatography and peptide mass was confirmed by electrospray ion trap mass spectrometry [27]. Myelin Basic Protein (MBP) was purchased from Enzo life sciences, antiMBP polyclonal antibody was obtained from Sigma Aldrich (St. Louis, MO, USA).

### 2.3. Induction and clinical evaluation of EAE

EAE was induced in the four strains of mice with an identical scheme by injecting 50 µg of myelin oligodendrocyte glycoprotein (MOG)35–55 peptide (1 mg/ml in saline) emulsified in IFA containing 4 mg/ml of M. tuberculosis. On day 0, after anaesthesia, we injected subcutaneously 50 µl of the emulsion in each of the two hind footpads and immediately thereafter 100 ng pertussis toxin in 50 µl saline was intravenously (i.v.) administered in the tail vein. On day 2, a second dosis of pertussis toxin was i.v. administered in the tail vein.

Mice were evaluated daily for signs of clinical disease with the following grading system: grade 0, normal; grade 0.5, floppy tail; grade 1, tail paralysis and mild impaired righting reflex; grade 2, mild hind limb weakness and impaired righting reflex; grade 3, moderate to severe hind limb paresis and/or mild forelimb weakness; grade 4, complete hind limb paralysis and/or moderate to severe forelimb weakness; grade 5, quadriplegia or moribund; grade 6, death.

### 2.4. Cell preparation from various organs

After euthanasia and perfusion of the mice, spleens were isolated, cut into small pieces and passed through cell strainers, to obtain single cell suspensions. Red blood cells were lysed by two incubations (5 and 3 min at 37°C) of the splenocyte suspensions in 0.83% NH_4_Cl solution. Remaining cells were washed two times with ice-cold PBS containing 2% FCS. For central nervous system (CNS) analysis, euthanized mice were gently perfused through the left cardiac ventricle with 50 ml ice-cold PBS to eliminate intravascular contaminating blood cells in the CNS. Spinal cords were removed by flushing the spinal canal with sterile PBS and brains were dissected. Brain and spinal cord cell fractions from individual mice were isolated according to the recent protocol by Legroux *et al.* [28]. Briefly, brains and spinal cords were digested with collagenase D and DNase for 15 min at 37^°^C. The products of the digestion were homogenized and filtered through a cell strainer (Becton Dickinson Labware, Franklin Lakes, NJ, USA) and centrifuged in a single 37% Percoll^TM^ step (10 min, 300 g, 4°C). The cell fractions were then washed twice by adding Hank’s Balanced Salt Solution (HBSS1X).

### 2.5. Flow cytometry analysis

Single cell suspensions (0.5 × 10 ^6^ cells) were incubated for 15 min with Fc-receptor-blocking antibodies anti-CD16/anti-CD32 (BD Biosciences Pharmingen, San Diego, CA, USA), washed with PBS supplemented with 2% FCS and then stained for 30 min with the indicated conjugated antibodies. Cells were washed twice and fixed with 0.37% formaldehyde in PBS. FITC-conjugated anti-CD19, PE-conjugated anti-CD3, APC-conjugated anti-CD4, BV410-conjugated anti-CD8, APC-conjugated anti-CD11b, BV711-conjugated anti-CD11c, FITC-conjugated anti-Gr-1 and PE-conjugated anti-F4/80, were purchased from eBioscience (San Diego, CA, USA). Cells were analysed in a FACS Fortessa flow cytometer and data were processed with the FlowJo software (Becton Dickinson Labware, Franklin Lakes, NJ, USA).

### 2.6. Zymography

Samples of tissue extracts were subjected to zymography analysis, as detailed previously. Briefly, to reduce background levels caused by contaminating glycoproteins, all samples were prepurified by gelatin-Sepharose affinity chromatography [29]. To allow quantitation and comparison of different substrate gels, all samples were spiked with a known amount of a recombinant deletion mutant of human proMMP-9 lacking the O-glycosylated and hemopexin domains, proMMP-9ΔOGHem [30]. The prepurified samples were loaded onto 7.5% polyacrylamide gels which contained 0.1% gelatin. After electrophoresis the gels were removed from the electrophoresis system and washed twice for 20 minutes with 2.5% Triton X-100. Then, the gels were incubated overnight at 37°C in 50 mM Tris-HCl pH 7.5, 10 mM CaCl_2_ for the development of gelatinolysis. Finally, the proteins in the gels were stained with Coomassie blue (GE Healthcare, Piscataway, NJ, USA) resulting in clear lysis zones which were analyzed with the ImageQuant TL software (GE Healthcare, Piscataway, NJ, USA). The concentrations of the gelatinase forms were calculated based on the density of the bands, on their relation with a known amount of a spiked internal standard sample of ΔOGHem proMMP-9 and on the comparison with a dilution series of a recombinant proMMP-9 standard mixture (including the ΔOGHem proMMP-9 that was used in the spiking) with known concentrations [31].

### 2.7. Multiplex ELISA

The protein levels of interferon (IFN)-α, IFN-γ interleukin (IL-6), tumor necrosis factor (TNF)-α, keratinocyte chemoattractant (KC) (or (GRO)-α or CXCL1), monocyte chemoattractant protein (MCP)-1 (or CCL2), MCP-3 (or CCL7), IP-10 (or CXCL10), macrophage inflammatory protein (MIP)-1α (or CCL3) and MIP-1β (or CCL4) were measured in plasma using a custom made 12 multiplex Assay (ProcartaPlex, eBioscience). The protocol from the manufacturer was followed and quantitative data acquisition was with two different plate readers which gave similar results: Luminex plate platform provided by the Laboratory of Virology, (Rega Institute for Medical Research, KU Leuven, Belgium) and with Magpix (ProDigest, University of Gent, Belgium).

### 2.8. Statistical analyses

Statistical analysis were performed using GraphPad Prism 6 software. Differences in the clinical course of EAE were analysed with a non-parametric Kruskal-Wallis test with Dunn’s multiple comparison. Significant differences between groups were evaluated using a non-parametric ANOVA Kruskal-Wallis test. All p values of 0.05 or less were considered significant.

## 3. Results

### 3.1. EAE develops in WT and in *mmp-9^−/−^*, but less in single *fas^lpr^* and *mmp-9^−/−^/fas* ^lpr^ mice

Previously, we documented the spontaneous disease phenotypes of single *fas*^lpr^, mmp-9^−/−^ mice and double *mmp-9^−/−^/fas^lpr^* mice, all on C57Bl/6j genetic background and without any exogenous (auto)antigenic stimulus. These studies generated the concept that MMP-9 plays on long-term a protective role in lupus-like syndromes [20]. Thereafter, we started to investigate whether this concept might be broadened to the prototypic organ-specific autoimmunity animal model of EAE. We induced CNS disease with MOG peptide and compared EAE development in wild-type (WT), single *fas^lpr^*, single *mmp-9*^−/−^ mice and to define the interactions between both Fas and MMP-9, we used double *mmp-9*^−/−^*/fas*^lpr^ mice [20]. Disease read-outs during the first five weeks corroborated previous findings with single *fas^lpr^* [32] and single *mmp-9* ^−/−^ mice [14]. These results were complemented with data about double *mmp-9^−/−^/fas^lpr^* mice (Figure 1A, 1B and 1C and Supplemental Table 1). In Figure 1A we illustrate, up to 35 days, that adult *mmp-9*^−/−^ mice developed EAE and even showed severed (P < 0.001 compared with WT mice) disease scores after about four weeks. These *mmp-9*^−/−^ mice also showed a slight delay in the disease onset (day 11), compared with the WT animals (day 10), (Figure 1C) as was described previously [14]. The fas^*lpr*^ mice were resistant against EAE development from two weeks onwards, as shown by Waldner et al. [32]. It is important to remark that although mmp-9^−/−^ animals showed a delay in the onset of the disease, they rather showed higher mean disease scores, thus the disease scores of the WT animals increased slowly while in the *mmp-9*^−/−^ cohort the disease appeared more aggressively, reaching higher disease scores. At day 10 after EAE induction, 70% of WT mice had already EAE symptoms, whereas in the cohort of mmp-9^−/−^ animals only 40% presented with symptoms, but the disease scores of the sick animals were higher in *mmp-9*^−/−^ animals than in WT mice at day 10. Data from the double *mmp-9^−/−^/fas^lpr^* mice hinted to independent actions of Fas and MMP-9. Whereas MMP-9 attenuated the lymphoproliferation syndrome, observed on long term in the double KO mice [20], it did not alter the EAE disease on the short term of 35 days (Figure 1A, 1B and 1C and supplemental table 1). Interestingly, *mmp-9^−/−^/fas^lpr^* mice also showed a delay in the disease onset (day 16) compared with the fas^*lpr*^ animals (day 13), (Figure 1C). In addition, lack of MMP-9 in the *fas^lpr^* mice (*mmp-9^−/−^/fas^lpr^*) resulted in a longer remission phase. *Fas^lpr^* mice were completely recovered from EAE symptoms at day 35, whereas in the *mmp-9^−/−^/fas^lpr^* 25 % of the animals still presented EAE symptoms (Figure 1A and B and supplemental table 1), suggesting different roles of MMP-9 in the initiation and the remission of the disease.

**Figure 1.**
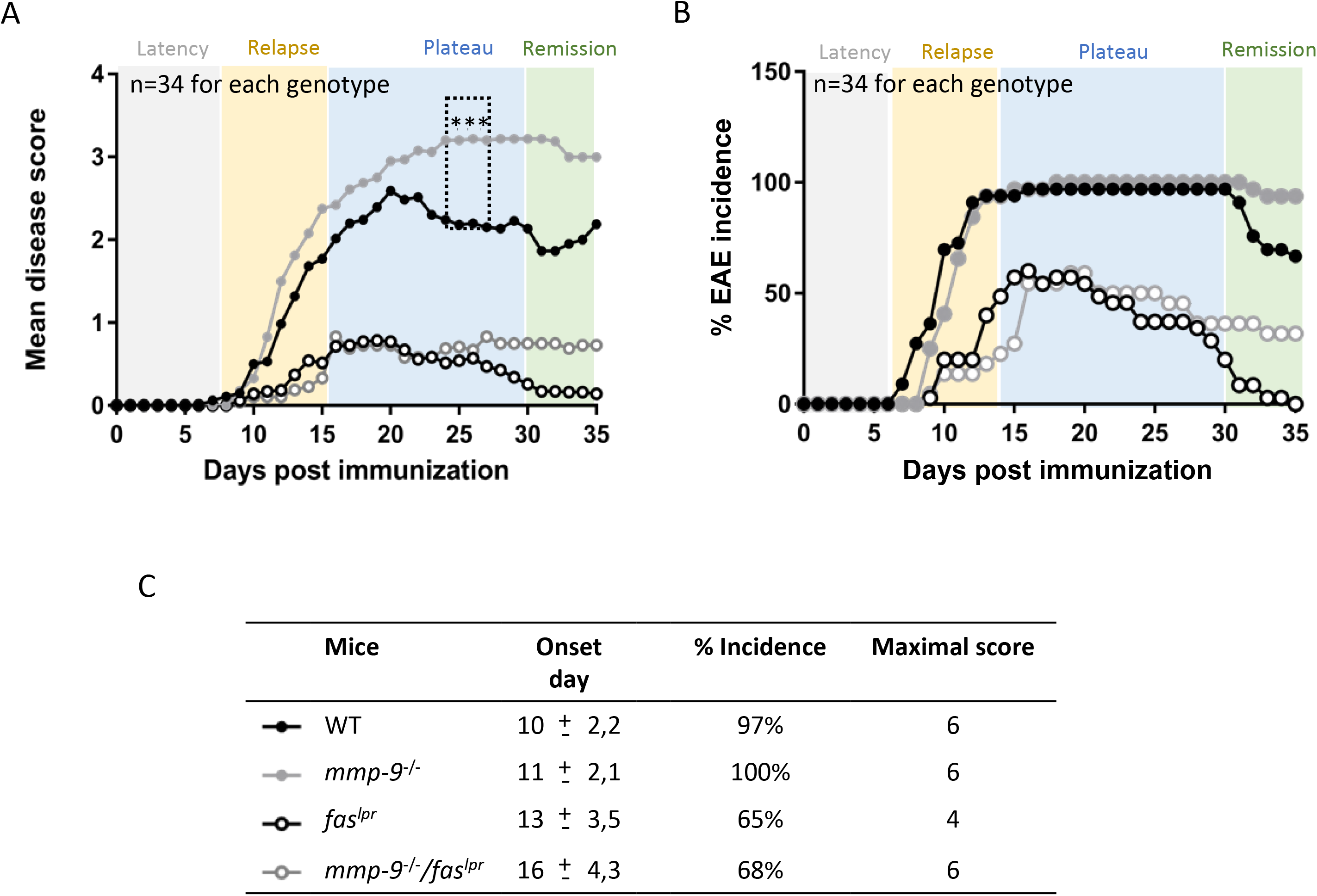
Clinical evolution of EAE in WT, *mmp-9*^−/−^, *fas*^lpr^ and *mmp-9^−/−^/fas*^lpr^ on short-term: EAE was induced in the four mouse strains and disease scores were measured during 35 days. (A) Mean disease scores from 34 mice for each of the four mouse strains. Significant differences in the comparison between WT and mmp-9^−/−^ are indicated with asterisks. Differences in mean diseases scores for all the data points from day 13 onwards till day 35 were significant for *fas^lpr^* and *mmp-9^−/−^/fas^lpr^ versus* WT. (B) Incidence of the disease after EAE induction. (C) Averages of day-of-onset, percentages of the incidence and maximal disease scores. (mean±SEM) ^*^p<0.05 *versus* WT by Kruskal-Wallis test with Dunn’s multiple comparison.

### 3.2. Alterations in peripheral blood and CNS leukocyte subsets in the EAE model (short-term)

With the knowledge that the infiltration of inflammatory cells into the brain parenchyma leads to clinical effects [17], we measured CNS cell populations after extensive systemic perfusion of the mice to eliminate blood leukocytosis effects. CNS cells were isolated from individual mice [28] at 1 month of the EAE experiment and characterized by flow cytometry to detect neutrophils, macrophages, dendritic cells, B lymphocytes, CD4-positive T helper and CD8-positive cytotoxic (Figure 2A). In general, the basal brain parenchymal cell pools were analogous in mice of the 4 genotypes. After EAE induction, all tested leukocyte subsets were significantly increased including in the central nervous system (CNS) for the four compared genotypes, including *fas^lpr^* and *mmp-9^−/−^fas^lpr^* mice, which presented lower incidence and diseases scores than WT and *mmp-9^−/−^* mice. Therefore, in these models no correlation exists between the clinical manifestations and the accumulation of the leukocytes into the CNS.

**Figure 2.**
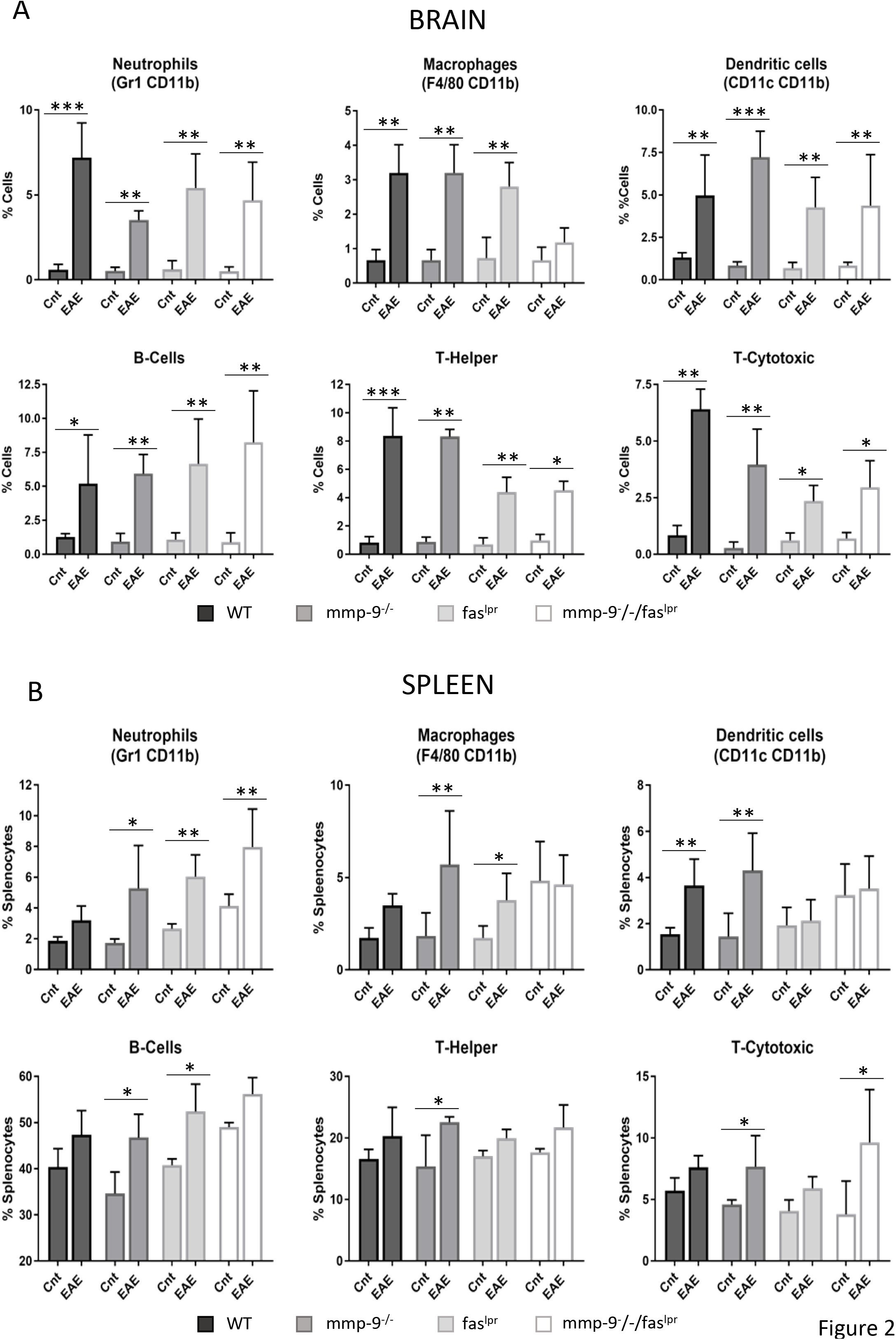
Induction and alterations of leukocyte subsets in CNS and spleens after 1 month of EAE initiation: (A)- Cells purified from spinal cord and brain were analyzed by flow cytometry with the use of the indicated markers. Neutrophils (Gr1^+^, CD11b^+^), macrophages (F4/80^+^, CD11b^+^), dendritic cells (CD11c^+^, CD11b^+^), B-cells (CD19^+^), T-helper cells (CD3^+^, CD4^+^) and T-cytotoxic cells (CD3^+^, CD8^+^) were analyzed in total tissue cell suspensions. (B)-Flow cytometry analysis of splenocytes for the indicated markers were used to identify neutrophils (Gr1^+^, CD11b^+^), macrophages (F4/80^+^, CD11b^+^), dendritic cells (CD11c^+^, CD11b^+^), T-helper cells (CD3^+^, CD4^+^), T-cytotoxic cells (CD3^+^, CD8^+^) and B-cells (CD19^+^). P values were determined by ANOVA Kruskal-Wallis test. ^*^p < 0.05, ^**^p<0.01 and ^***^p<0.001. The histogram bars represent the average of all the data points. Cnt: Control.

In addition, we monitored spleen leukocyte subsets in the ongoing plateau (1 month) of inflammation. In Figure 2B, the percentages of the counts of neutrophils, macrophages, dendritic cells, B lymphocytes, CD4-positive T helper and CD8-positive T cytotoxic cells in spleens are represented. No mayor differences were observed between the four different genotype groups. In general, at 1 month after EAE induction, the relative abundancies of all subsets of splenic leukocytes were slightly increased. In particular, all the splenic leukocyte studied were significantly increased by EAE induction in the *mmp-9*^−/−^ mice. In addition, neutrophils were significantly increased in *fas^lpr^* and double *mmp-9* ^−/−^*/fas^lpr^* mice and macrophages and B lymphocytes in *fas^lpr^.*

### 3.3. Altered levels of cytokines and chemokines after short-term of EAE induction

Cytokines,chemokines and their receptors play major roles in de progression of MS and EAE [12]. Therefore, we analyzed the plasma levels of the chemokines: GRO-alpha/KC (or CXCL1), MCP-1 (or CCL2), MCP-3 (or CCL7), IP-10 (or CXCL10) and MIP-1α (or CCL3) (Figure 3A) and the cytokines TNF-α, IL-6, IFN-α and IFN-γ (Figure 3B). The levels of many of the selected cytokines and chemokines were increased after 1 month of EAE induction but not all increases were found to be significant. A significant increase of TNF-α was consistently detected in mice of all four genotypes 1 month after EAE induction. IFN-α, KC, MIP-1α and MCP-1 levels were significantly increased in *fas^lpr^* mice. The levels of IP-10 and MIP-1α were increased significantly in WT, *mmp-9*^−/−^ and *fas^lpr^*, but not in the double *mmp-9^−/−^/fas^lpr^* mice. IL-6 levels were only significantly increased in WT and mmp-9^−/−^ animals. In the case of fas^*lpr*^ and double *mmp-9^−/−^/fas^lpr^* mice, IL-6 levels had only a trend towards increases at 1 month after EAE induction.

**Figure 3.**
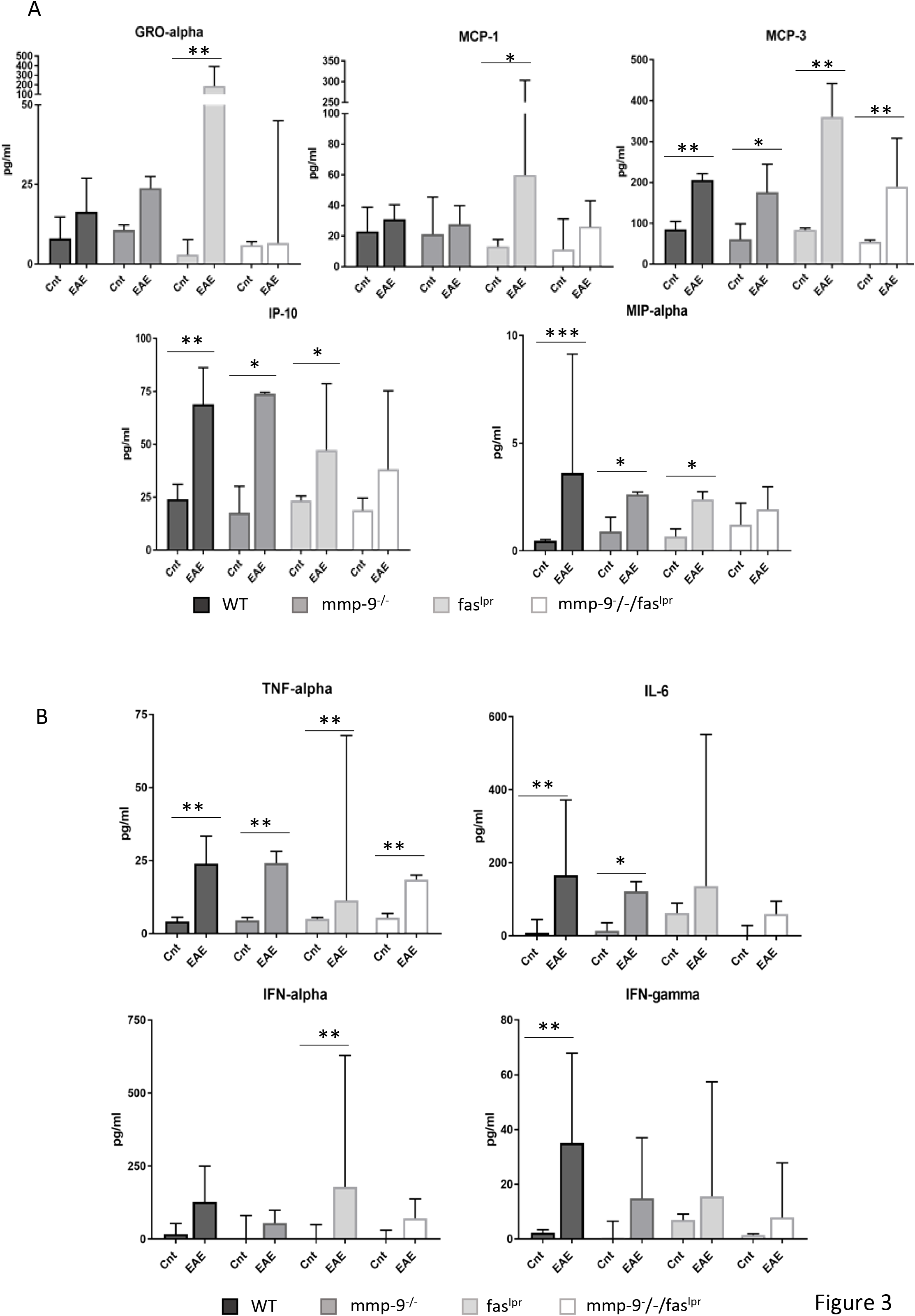
Plasma chemokine and cytokine levels at 1 month after EAE induction: ELISA analysis of the indicated chemokines (A) and cytokines (B) from plasma samples obtained after 1 month in control animals (Cnt) and animals with EAE (EAE). The histograms represent the average of each group. P values were determined by ANOVA Kruskal-Wallis test. ^*^p<0.05, ^**^p*0.01 and ^***^p<0.001 *versus* control 1 month.

### 3.4. Long-term EAE in WT, single *mmp-9^−/−^*, *fas^lpr^* and double *mmp-9^−/−^/fas*^lpr^ knockout mice

In a second type of experiment, we analyzed long-term evolution of EAE in smaller animal cohorts, and we compared the clinical evolutions in mice of the four genotypes up to 100 days, allowing all the animals to reach complete remission. As shown in Figure 4 all animals had complete clinical recovery after maximally 47 days. In the evaluations of the four disease phases (latency, relapse, plateau and remission), we remarked that *mmp-9^−/−^* and *mmp-9^−/−^/fas^lpr^* mice animals had again a slightly later onset of the disease than their respective controls WT and *fas^lpr^* mice (day 10, 11 for WT and *mmp-9*^−/−^, and day 13, 15 for *fas^lpr^* and *mmp-9^−/−^/fas^lpr^* mice, respectively). Interestingly, the remission phase was significantly extended in the *mmp-9^−/−^* cohort, which needed more time to completely recover than the wild type animals (10 days more than WT mice). This data were suggestive for roles of MMP-9 in auto-antigen production (latency and relapse phases) and clearance of autoantigens and immune complexes (plateau and remission phases), the latter being as previously suggested for the SLE Fas/MMP-9 model [20].

**Figure 4.**
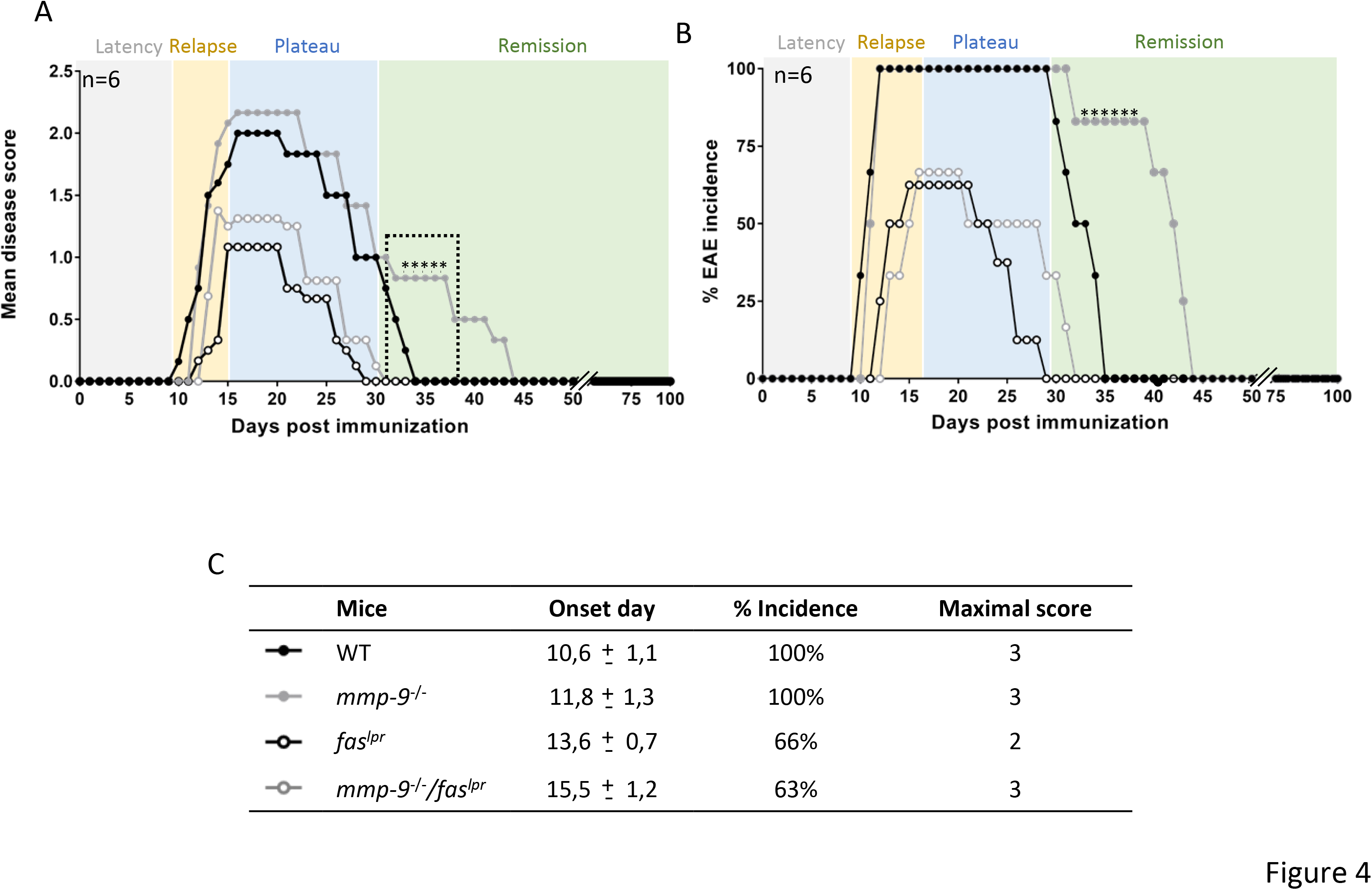
Clinical follow-up of EAE in WT, *mmp-9*^−/−^, faslpr and *mmp-9*^−/−^*/fas*^lpr^ for 100 days: EAE was induced in groups (n=6) of mice of the four different genotypes and disease scores were measured during 100 days. (A) Significant differences between WT and mmp-9−/− are indicated with asterisks. (B) Percentages of EAE incidence after the induction of the disease. (C) Average day-of-onset, percentage of incidence and maximal score. (mean±SEM) ^*^p<0.05 versus WT by Kruskal-Wallis test with Dunn’s multiple comparison.

### 3.5. Alterations in CNS and spleen leukocyte subsets in the EAE long-term model (100 days)

We then studied the CNS and spleen leukocyte subsets in the fading out of inflammation. Significant CNS inflammation was not observed anymore at 3 months. Nevertheless, there was still a slightly increase in the levels of neutrophils and T-cytotoxic cells in the EAE induced mice from all the 4 genotypes studied compared with the controls (Figure 5A), which indicated that inflammation in the CNS was still present. Only significant increased numbers of cytotoxic T lymphocytes in the CNS of the single *mmp-9*^−/−^ mice at 3 months was observed. This observation is in line with the prolonged disease phase before remission (Figure 4 A and B) in the *mmp-9*^−/−^ mice in comparison with WT mouse cohort.

**Figure 5.**
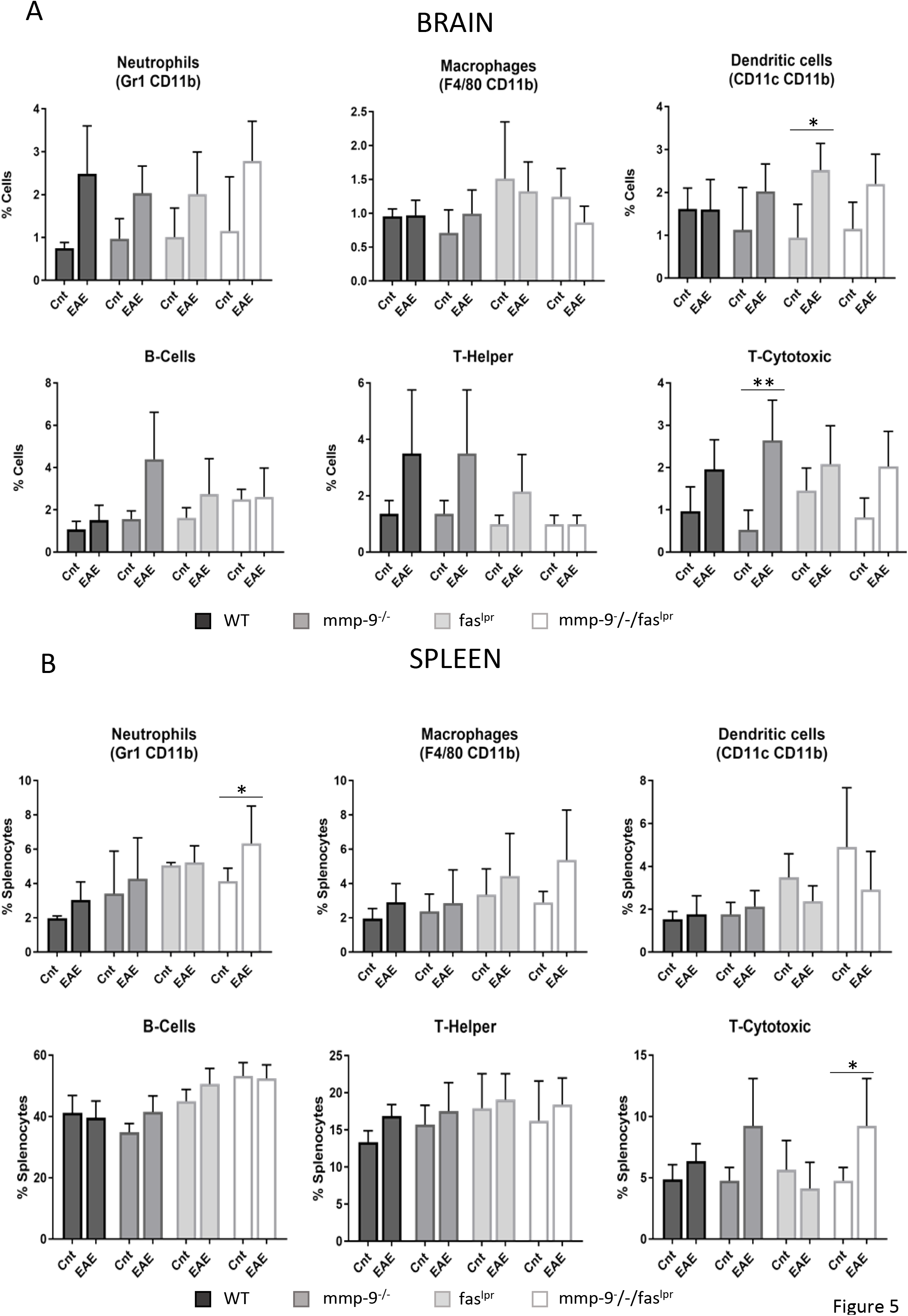
Changes in leukocyte percentages in the CNS and spleens of mice with EAE after 100 days: Cells purified from spinal cord and brain (A) and spleens (B) were analyzed by flow cytometry with the use of the indicated markers. Neutrophils (Gr1^+^, CD11b^+^), macrophages (F4/80^+^, CD11b^+^), dendritic cells (CD11c^+^, CD11b^+^), B-cells (CD19^+^), T-helper cells (CD3^+^, CD4^+^) and T-cytotoxic cells (CD3^+^, CD8^+^) were analyzed in total tissue cell suspensions. P values were determined by ANOVA Kruskal-Wallis test. ^*^p<0.05 and ^**^p<0.01. The histogram bars represent the average of all the data points. Cnt: Control.

At the remission phase, splenic leukocyte populations in the WT mice and both single KO mice had returned to control levels (Figure 5B). However, neutrophils and CD8 T cells levels remained significantly increased in the *mmp-9^−/−^/fas^lpr^* mice.

### 3.6. Altered levels of cytokines and chemokines in a long-term after EAE induction

Next we studied the levels of chemokines (Figure 6A) and cytokines (Figure 6B) after recovery at 100 days after EAE induction, thus in the absence of EAE symptoms. Interestingly, although the levels of most of the cytokines and chemokines decreased to control levels after 3 months of EAE induction, TNF-α and MIP-1α and IP-10 still remained significantly higher in comparison with control healthy animals for the WT and single mmp-9^−/−^ mice, but not for the double fas^lpr^ and *the mmp-9^−/−^/fas^lpr^* mice.

**Figure 6.**
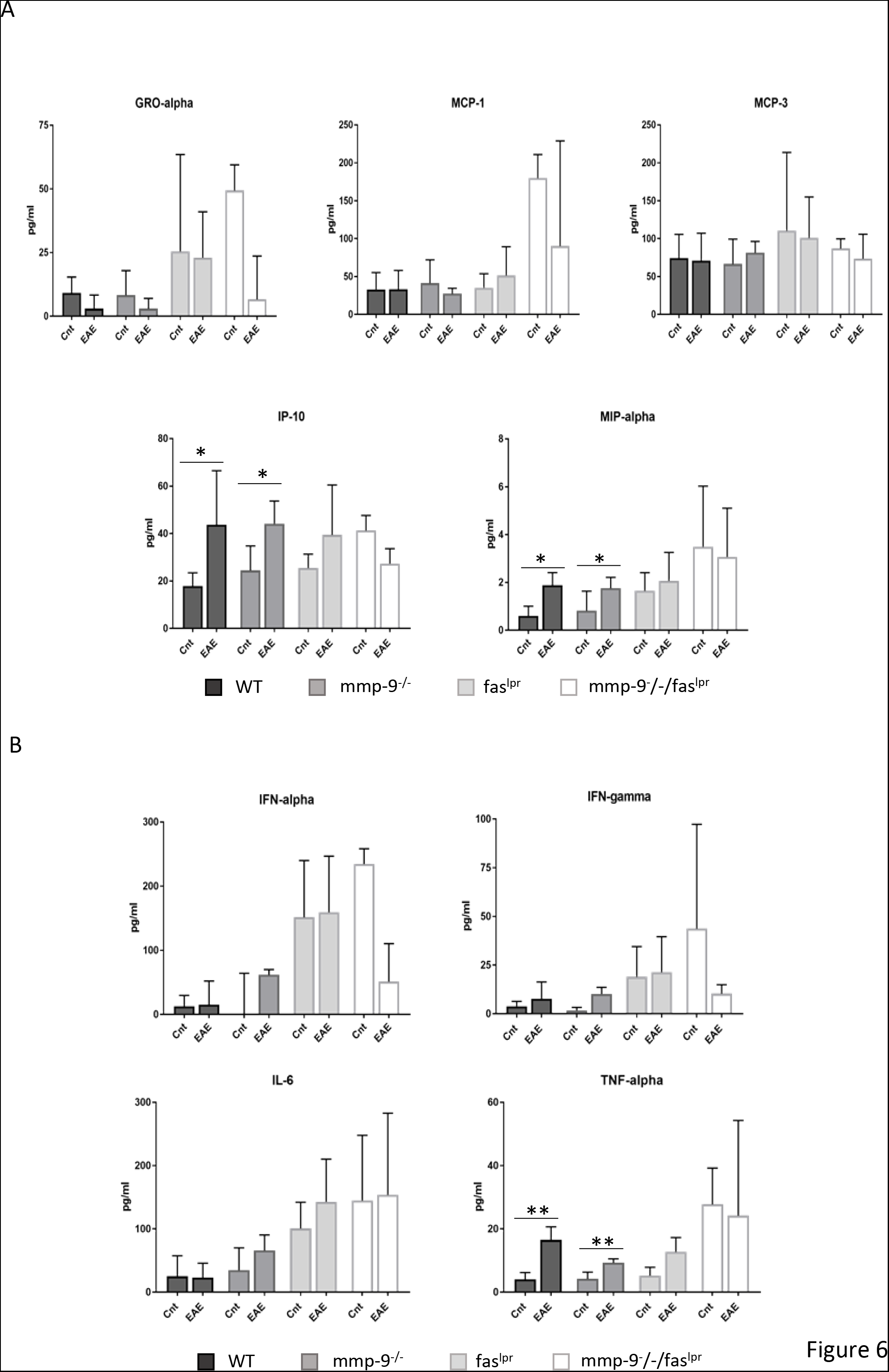
Plasma chemokine and cytokine levels at 100 days after EAE induction: ELISA analysis of the indicated chemokines (A) and cytokines (B) from plasma samples obtained after 100 days in control animals (Cnt) and animals with EAE (EAE). The histograms represent the average of each group. P values were determined by ANOVA Kruskal-Wallis test. ^*^p<0.05 versus control.

### 3.7. EAE phase-specific changes in MMP-9 in WT and *fas^lpr^* mice

Because MMP-9 is a pivotal marker of inflammation in the CNS [12,17,33] and *mmp-2/mmp-9*-double knockout mice are resistant against EAE development [15,17], we evaluated the expression levels of the gelatinases MMP-2 and MMP-9 in the CNS and spleen (as a peripheral tissue) with optimized zymography analysis [31]. As previously demonstrated, by inducing EAE, MMP-9 levels were enhanced significantly in the CNS at 1 month (Figure 7A). Unexpectedly, in the immunized *fas^lpr^* mice the MMP-9 levels in the brain were also significantly increased (Figure 7A), whereas these animals presented with significant lower disease scores than WT mice. Furthermore, the levels of MMP-2 did not change significantly, neither by induction of EAE, nor by deletion of *fas* or *mmp-9* (Supplementary Figure 1). This is in line with the view that MMP-2 is generally a constitutive enzyme in inflammation [12,24,34]. As a complementation, we evaluated MMP-9 tissue levels in the spleen as read-out of systemic organ effects (Figure 7B). MMP-9 and interestingly activated MMP-9 levels were increased significantly, at 1 month, by EAE induction in WT and in *fas* ^lpr^ mice.

**Figure 7.**
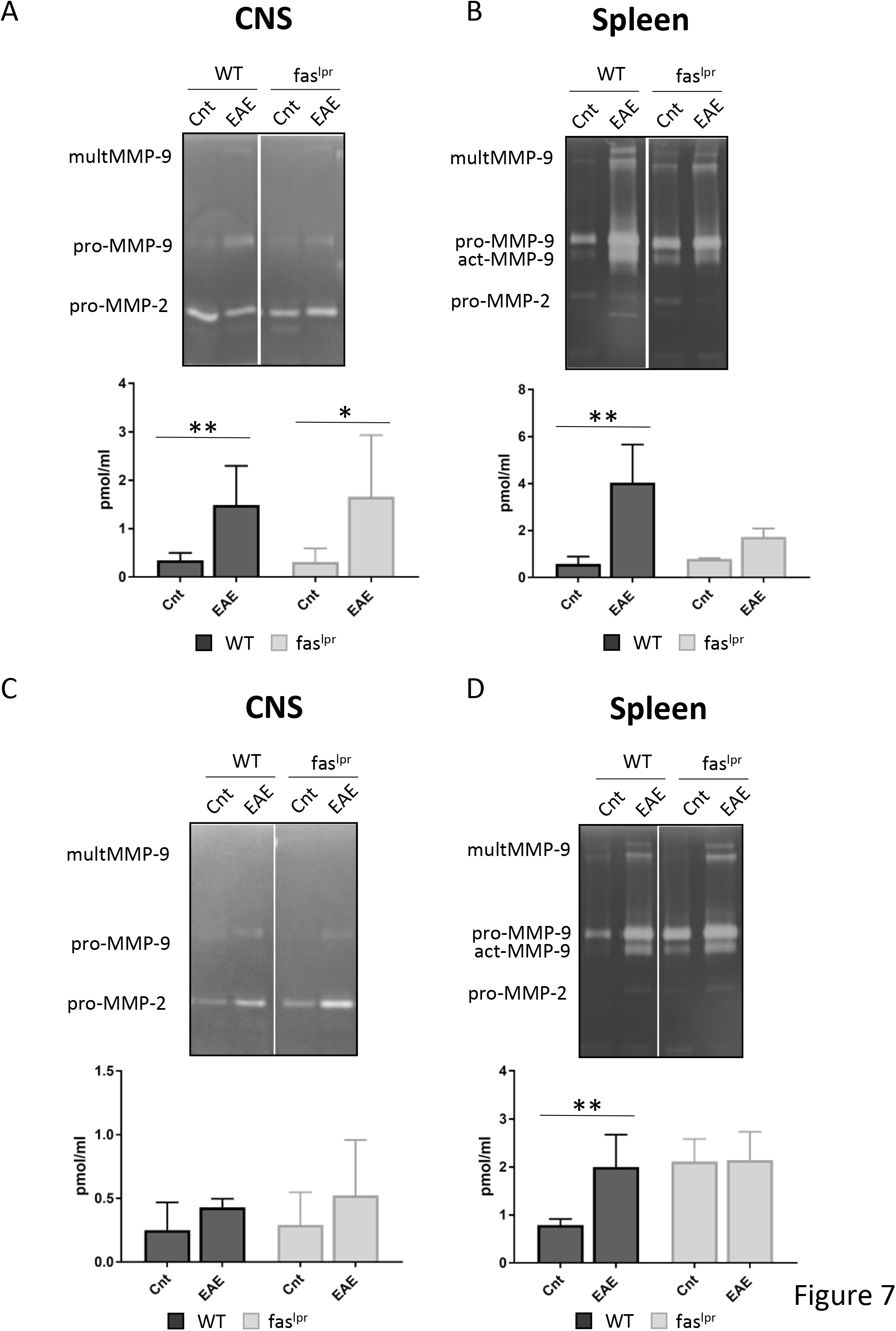
MMP-9 upregulation in animals with EAE: Quantitative gelatin zymography analysis was used to analyze MMP-9 on tissue extracts from the central nervous system and spleens after 1 month (A) and 100 days (B) of EAE induction. Volumes equivalent to 40 µl of tissue extracts were prepurified by gelatin-Sepharose. After purification, the samples were analyzed by gelatin zymography (gels) and the bands were quantified with the use of in-depth standardization. The histograms represent averages of the quantification of MMP-9 per. P values were determined by ANOVA Kruskal-Wallis test. ^*^p<0.05 and ^* *^p<0.01. Cnt 1m: Control at 1 month. 1m: EAE at 1 month. Cnt 3m: Control at 3 months. 3m: EAE at 3 months.

We also analyzed and compared MMP-9 levels at 3 months after EAE induction (Figure 7C and D). In this analysis, we observed a decrease to basal levels locally in the CNS, both for WT and *fas*^lpr^ mice (Figure 7C). In contrast, MMP-9 levels remained high in the spleen of WT mice (Figure 5B). Although the WT animals had no longer signs of illness at that moment, the obtained data indicated that peripheral inflammation was still present.

### 3.8. MBP is degraded in immune complexes

*Mmp-9^−/−^* animals showed increased disease scores and a delayed recovery in comparison with WT mice, thus suggesting that MMP-9 might play different roles in the pathogenesis of EAE, depending on an early or late disease phase [19]. Previously we showed that MMP-9 cleaves free MBP [35,36] and thus generates remnant epitopes of this auto-antigen, whereas independently and on long term MMP-9 might also contribute to the clearance of autoantigens [20]. We hypothesized that MMP-9 might clear immunogenic MBP peptides as such, but also might cleave MBP peptides caught within immune complexes (IC) between anti-MBP and MBP. Therefore, we analyzed the cleavage pattern of MBP in free form and in the form of IC. In Figure 8 it is shown that MBP was cleaved similarly when it was free or present in IC. These data are the first to indicate that MMP-9 is capable to destroy not only free autoantigens but also such autoantigens captured within IC.

**Figure 8.**
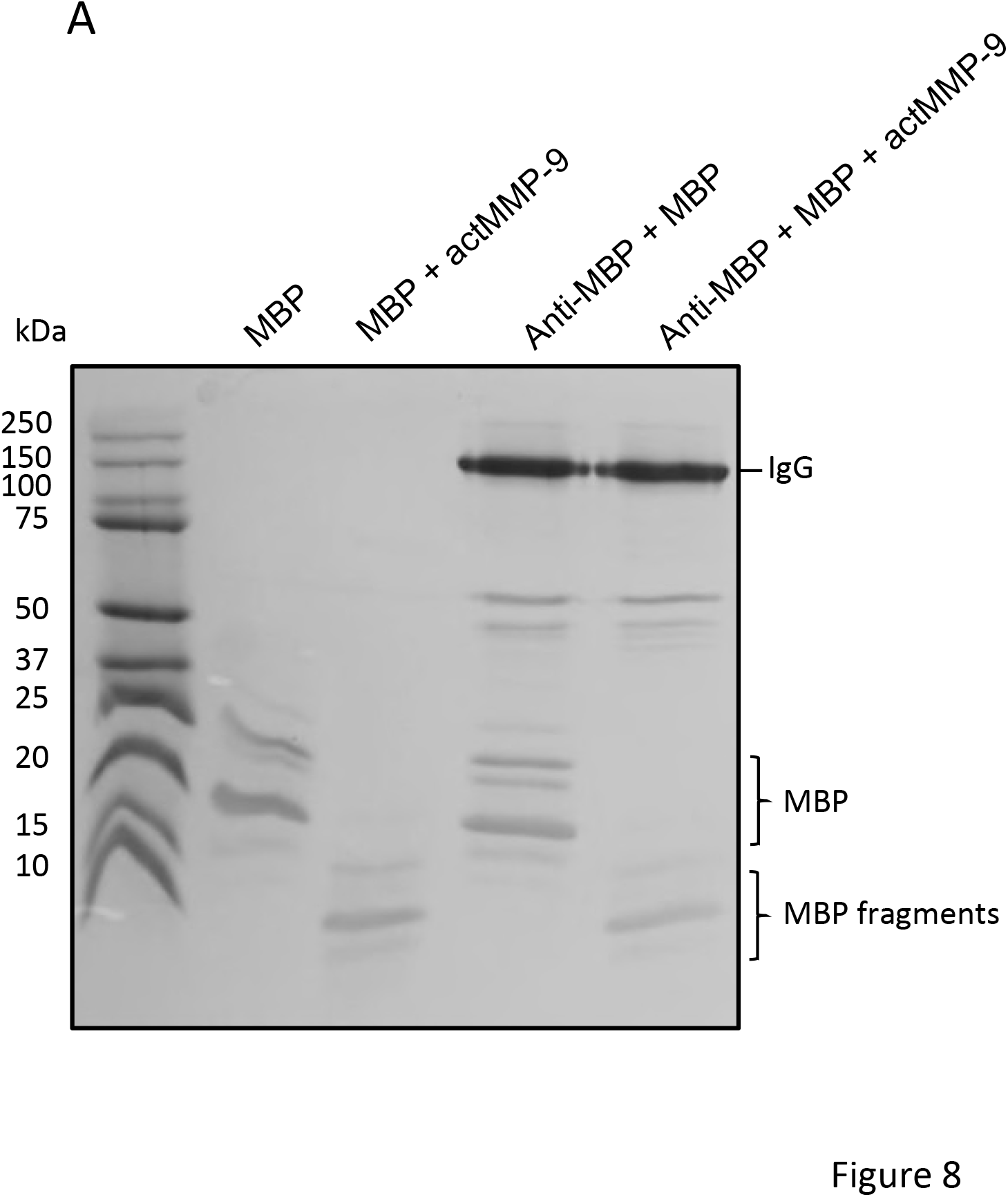
MBP and anti-MBP analysis: (A) Aliquots of 2 µg of MBP were incubated with anti-MBP antibody in an excess 1:4 molar ratio MBP:Ab during 15 minutes. Next, active MMP-9 was added at an enzyme:substrate ratio of 1:1000. After 16h, the reaction products of the incubations were analyzed by non-reducing SDS-PAGE and Coomassie blue staining. Immunoglobulin G (IgG), myelin basic protein isoforms (MBP) and MBP fragments are indicated.

## 4. Discussion

Thanks to the use of preclinical studies with gene knockout mice and protease inhibitors, it has been suggested that MMPs, and MMP-9 in particular, may be therapeutic targets in MS. Although young *mmp-9^−/−^* mice are protected against EAE, adult *mmp-9^−/−^* mice develop EAE [14]. Therefore, an aggravating role for MMPs has been documented in EAE [15]. MMP-2 and other proteases in an inflammatory network [37,38] may act together to establish or control clinical disease, which can eventually be reversed with MMP inhibitors. An extreme phenotype was the observation of complete protection against EAE in young *mmp-2/mmp-9* double knockout mice [15]. However, in most preclinical studies only short-term effects of MMP-9 gene deletion or protein inhibition on EAE have been studied, therefore we performed a long-term EAE study to analyze the role of MMP-9 after clinical remission.

MMP-9 plays a protective role in systemic autoimmunity in vivo, induced by fas deficiency [20] and single *fas^lpr^* mice are protected against EAE [8–10, 32]. Whether the induction of EAE is extinguished or aggravated in double *mmp-9^−/−^/fas^lpr^* knockout mice was not known. For that reason, we crossed *fas* ^lpr^ *mice, prone to systemic autoimmunity, with mmp-9* ^−/−^ mice to dissect the net effect of MMP-9 on EAE development on short- and long-terms in an antigen-driven model of organ-specific autoimmunity.

In short-term EAE experiments, we showed that *mmp-9^−/−^* mice presented a slight delay in the disease onset but higher disease scores compared with WT mice, whereas, as expected, fas^*lpr*^ andc *mmp-9^−/−^/fas^lpr^* mice had minor disease symptoms and lower disease incidence. Interestingly, infiltration of leukocytes in the CNS and cytokine and chemokine levels were increased in the four genotypes studied, independently of the disease scores.

Analysis of the long-term EAE experiments, suggested a protective role of MMP-9 in the remission phase of the disease, thus *mmp-9^−/−^* mice needed longer time intervals to completely recover from the EAE symptoms. Furthermore, TNF-α, MIP-1α and IP-10 levels remained significantly higher in the WT and *mmp-9*^−/−^ mice after 100 days of the EAE induction.

Due to the role of activated MMP-9 in breaking the blood brain barrier in MS [12,17], we expected a protective effect of deletion of MMP-9 on the induction of EAE in adult mice. In line with this expectation, disease onset was slightly delayed in *mmp-9* ^−/−^ mice, but in contrast, we observed in adult mice a significantly increased disease score during the plateau phase (Figure 1), which suggested that MMP-9 acted as a protective factor once the injury is made. This idea was reinforced by the observation of a delay in the remission of mmp-9^*-*^/- mice in comparison with WT mice (Figure 4A and B).

MMP-9 processes chemokines [39] plays an important role in cytokine and chemokine regulation in many pathologies and animal models of disease, including EAE. MMP-9 and MMP-2 selectively cleave chemokines and modulate their activity or availability [39,40]. These proteases are also able to promote chemokine secretion by astrocytes after cleavage of Notch, which results in the inhibition of PTEN and the activation of the Akt/NF-kB pathway [41]. Such processes provide additional explanations why *mmp-9*^−/−^ animals present a slight delay in the onset of the disease. The lack of MMP-9 will delay the expression of chemokines by astrocytes and, as a consequence, the migration of immune cells to the brain. Contrarily MMP-9 could act as an inactivator of chemokines, for example, MMP-9 is able to cleave stromal-derived factor-1 (SDF-1/CXCL12), thereby inactivating this particular chemokine [42]. In the case of MCP-1/CCL2 and MIP-1a/CCL3, the fragment generated after cleavage by MMP-2 behaves as a chemokine receptor antagonist [43]. The absence of MMP-9 will preserve the levels of intact chemokines and delay the inactivation of these specific chemokines and consequently maintain the disease course in the MMP-9 knockout mice, once it is initiated. These mechanisms could explain why *mmp-9^−/−^* mice presented a significantly prolonged remission phase compared with WT controls.

Why *mmp-9*^−/−^ mice developed delayed but more exacerbated early EAE might also be explained by the overexpression of other MMPs [44]. For instance, Esparza and colleagues showed increased EAE in *mmp-2-KO* mice due to increased MMP-9 levels [45]. We tested the protein levels of MMP-2 in our *mmp-9*^−/−^ mice and these were not significantly altered. In addition, by RNAseq analysis we found that basal levels of all mouse MMP mRNAs, except that of MMP-9, were not altered in our WT (n=8) versus *mmp-9*-KO mice (n=8) [26].

On long term, MMP-9 may also have beneficial effects, for example in the remyelination process [46] and it protects against systemic autoimmunity in *fas*^lpr^ mice [20]. For these reasons it was imperative to investigate what happens in an (auto-)antigen-driven condition in the presence or absence of MMP-9.

We hypothesized that MMP-9 helped in the clearance of (auto)antigens and immune complexes (IC). This hypothesis is supported by the in vitro experiments in which we show how MMP-9 is able to degrade MBP, even when MBP is in immune complexed form together with antiMBP-Ab (Figure 8). We choose on purpose to study MBP and not MOG for two reasons: (i) because we induced EAE with MOG35–55 peptide, the adaptive immune response might be skewed towards this peptide but not necessarily towards MBP and (ii) MBP is a water-soluble substrate of MMP-9 for which the remnant epitopes are well studied [35, 44].

Although the processing of MBP was discovered with MMP-9 as a prototypic MMP [35], also MMP-2, MMP-8, MMP-10, MMP-12, MT1-MMP and MT6-MMP are MMPs able to process MBP and its proteolysis was superior with MT6-MMP [44]. Several fragments of MBP including peptides 1–15, 68–86, 83–99, 84–104 and 87–99 are known autoantigens. Although MMP-9 is capable to generate the immunogenic peptide MBP_1–15_, it is also able to cleave at residues 91, 93, 97, 102, 109, 134, and 153, thus contributing to the degradation and clearance of the immunogenic peptides MBP83–99, MBP84–104 and MBP87–99. One might ask whether the generation of MBP remnant epitopes [11] becomes a next stimulus to induce a new autoimmune attack. Whereas this is perfectly possible and can not be excluded, it must be recognized that the autoantigen clearance remains an independent protective function. Indeed, those antibodies that capture the remnant epitopes autoantigens with the highest affinities will opsonize and eliminate these maximally, because the IgG remained intact, even after prolonged cleavage by MMP-9 (Figure 8). To understand the pathophysiology of and to better treat MS and other autoimmune diseases, it is thus critical to distinguish between the (early) phase of autoantigen generation, the awakening phase of the adaptive immune response and, as we here further document, the plateau and resolution phases [19]. By our finding of long-term persistence of innate cytokine and protease responses, we advocate that, for evaluations of novel treatments in preclinical settings, a long-term monitoring of cytokines, chemokines and MMP-9 will discriminate better between curative and symptomatic treatments than the earlier EAE studies, done with observations for only limited duration.

Another critical aspect, gaining more attention in recent research on MS and EAE, is the involvement of B-lymphocytes and antibodies. The oligoclonal antibody response in the cerebrospinal fluid has been used for a long time as a pathognomonic diagnostic test for MS, although the role of these immunoglobulins remains enigmatic. Nevertheless, antigen antibody complexes are pro-inflammatory by, for instance, activating the classical complement pathway. Destruction and resolution of immune complexes in the late phase of autoimmunity by proteolysis may thus be anti-inflammatory and beneficial.

Brändle *et al.* proved that distinct oligoclonal band antibodies in MS recognize ubiquitous intracellular self-proteins not specific to the brain, suggesting that the B-cell responses may be partially directed against intracellular autoantigen release during tissue destruction [47]. The oligoclonal antibodies that target intracellular proteins might be the result of a secondary immune reaction against cellular debris. These reactivities could constitute a more general characteristic of autoimmune and inflammatory diseases instead of being specific for MS. This viewpoint was already defined long ago by Grabar [48]. MMP-9 is able to degrade a broad range of intracellular proteins [49], thus the lack of this gelatinase might affect the amounts and persistence of intracellular antigens, thus increasing the levels of oligoclonal antibodies against these persistent proteins. This might explain why *mmp-9*^−/−^ animals show higher disease scores, a prolonged plateau phase and a delayed remission in comparison with wild-type mice.

Mechanistically, the protective function of MMP-9 in the remission phase might be by autoantigen destruction and clearance, even for formed immune complexes. Whereas MMP-9 may be detrimental in the early phase of MS and EAE, once immune complexes with autoantigens have formed, MMP-9 becomes a protective factor by eliminating free and immune-complexed autoantigens.

As it was originally described, animals that lack FAS protein are protected to develop EAE [8–10, 32]. Surprisingly, the abundancies of CD4 and CD8-positive T cells, B cells, neutrophils, dendritic cells and macrophages were increased in the spleens and the central nervous system of these animals, even in these animals that did not show any disease symptoms. In addition, the MMP-9 levels in the brain and the spleen were increased suggesting that inflammation is present in these organs. The fact that these animals did not suffer from disease might be explained by lack of CNS cell apoptosis due the lack of FAS and disruption of the known neuronal apoptosis mechanism through Fas/Fas-ligand interaction [50,51]. A second explanation is the observation that only CNS leukocytes, which enter the brain parenchyma, cause disease symptoms. Leukocytes that are retained into vascular cuffs – which may become gigantic – do not cause disease symptoms [17]. It is only when the blood-brain barrier is destroyed and the leukocytes penetrate through the barrier into the brain parenchyma that clinical neurological signs become evident [41]. This also explains why, irrespective of the persistence of significantly more cytotoxic T lymphocytes in the brains and spinal cords at 3 months in the single *mmp-9*^−/−^ mice, no clinical disease was observed anymore.

MS patients show high cytokine levels even when they are in clinical remission phase, which implies that, although an association exists between specific cytokine profiles and the progression of the disease activity, MS patients have a constant and complex activation of the immune system [52]. Our data from long-term EAE studies were in line with this observation. We showed that the levels of % 8 Q IP-10, and ˙,’ Q remained significantly above basal levels in the groups with higher diseases scores (WT and *mmp-9^−/−^* mice) at 100 days after EAE induction, although the animals did not present anymore EAE symptoms. The latter situation may be correlated with high cytokine levels in the MS remission phase. Thus, although more studies need to be done, we suggest (i) that long-term studies after EAE induction represent a useful model to study the remission phase of MS patients and (ii) that therapy efficiency may be monitored with the analysis of such cytokines and chemokines, both preclinically and clinically.

**In conclusion, whereas Fas is a strong driver molecule of EAE, MMP-9 is an independent and phase-specific effector molecule. In the plateau and early induction phase of EAE MMP-9 contributes to the generation of remnant epitopes and initiation of adaptive immune processes. In the later remission phase, MMP-9 becomes a protective factor when autoantigens in either free form or immune-complex form need to be eliminated and cytokines inactivated.**

## 5. Acknowledgements

The present study was possible thanks to funding by the Geconcerteerde OnderzoeksActies (GOA 2013/015) and (C16/17/010), the Foundation for Scientific Research of Flanders (FWO-Vlaanderen) and the Charcot Foundation (Belgium).

## 6. Author contributions

Designed the study: G.O. and E.U.B. Performed experiments and analyzed data: E.U.B., N.B., L.B., E.M., J.V. P.P., B.C. J.V.D., and G. T. Generated, tested and kept the knockout mouse colonies: G.O. and G.T. Wrote the manuscript: E.U.B. and G.O. with input from all authors.

## 7. Additional Information

Competing financial interests: The authors declare no competing financial interests.

---

**Supplemental Figure 1. Zymography analysis from brain and spleen of mmp-9^−/−^ and mmp-9^−/−^fas^lpr^ mice:** Tissue extracts from CNS (A) and spleen (B) were analyzed by gelatin zymography. Cnt 1m: Control at 1 month. 1m: EAE at 1 month. Cnt 3m: Control at 3 months. 3m: EAE at 3 months.

